# CoronaHiT: High throughput sequencing of SARS-CoV-2 genomes

**DOI:** 10.1101/2020.06.24.162156

**Authors:** Dave J. Baker, Alp Aydin, Thanh Le-Viet, Gemma L. Kay, Steven Rudder, Leonardo de Oliveira Martins, Ana P. Tedim, Anastasia Kolyva, Maria Diaz, Nabil-Fareed Alikhan, Lizzie Meadows, Andrew Bell, Ana Victoria Gutierrez, Alexander J. Trotter, Nicholas M. Thomson, Rachel Gilroy, Luke Griffith, Evelien M. Adriaenssens, Rachael Stanley, Ian G. Charles, Ngozi Elumogo, John Wain, Reenesh Prakash, Emma Meader, Alison E. Mather, Mark A. Webber, Samir Dervisevic, Andrew J. Page, Justin O’Grady

**Author notes:** Corresponding authors (bioinformatics); (sequencing). Contributed equally (Dave J. Baker, Alp Aydin, Andrew J. Page, Justin O’Grady).

## Abstract

The COVID-19 pandemic has spread to almost every country in the world since it started in China in late 2019. Controlling the pandemic requires a multifaceted approach including whole genome sequencing to support public health interventions at local and national levels. One of the most widely used methods for sequencing is the ARTIC protocol, a tiling PCR approach followed by Oxford Nanopore sequencing (ONT) of up to 96 samples at a time. There is a need, however, for a flexible, platform agnostic, method that can provide multiple throughput options depending on changing requirements as the pandemic peaks and troughs. Here we present CoronaHiT, a method capable of multiplexing up to 96 small genomes on a single MinION flowcell or >384 genomes on Illumina NextSeq, using transposase mediated addition of adapters and PCR based addition of barcodes to ARTIC PCR products. We demonstrate the method by sequencing 95 and 59 SARS-CoV-2 genomes for routine and rapid outbreak response runs, respectively, on Nanopore and Illumina platforms and compare to the standard ARTIC LoCost nanopore method. Of the 154 samples sequenced using the three approaches, genomes with ≥ 90% coverage (GISAID criteria) were generated for 64.3% of samples for ARTIC LoCost, 71.4% for CoronaHiT-ONT, and 76.6% for CoronaHiT-Illumina and have almost identical clustering on a maximum likelihood tree. In conclusion, we demonstrate that CoronaHiT can multiplex up to 96 SARS-CoV-2 genomes per MinION flowcell and that Illumina sequencing can be performed on the same libraries, which will allow significantly higher throughput. CoronaHiT provides increased coverage for higher Ct samples, thereby increasing the number of high quality genomes that pass the GISAID QC threshold. This protocol will aid the rapid expansion of SARS-CoV-2 genome sequencing globally, to help control the pandemic.

## Introduction

The COVID-19 pandemic caused by the SARS-CoV-2 virus began late 2019 in Wuhan, China and has now spread to virtually every country in the world, with tens of millions of confirmed cases and millions of deaths (Dong, Du, and Gardner 2020). Key to the control of the pandemic is understanding the epidemiological spread of the virus at global, national and local scales (Shu and McCauley 2017). Whole genome sequencing of SARS-CoV-2 is likely to be the fastest and most accurate method to study virus epidemiology as it spreads. We are sequencing SARS-CoV-2 as part of the COVID-19 Genomics UK (COG-UK) consortium, a network of academic and public health institutions across the UK brought together to collect, sequence and analyse whole genomes to fully understand the transmission and evolution of this virus (https://www.cogconsortium.uk/). The SARS-CoV-2 genome was first sequenced in China using a metatranscriptomic approach (Wu et al. 2020). This facilitated the design of tiling PCR approaches for genome sequencing, the most widely used of which is the ARTIC Network (https://artic.network) protocol. Consensus genome sequences are typically made publicly available on GISAID (Elbe and Buckland_JMerrett 2017). This has enabled real-time public health surveillance of the spread and evolution of the pandemic through interactive tools such as NextStrain (Hadfield et al. 2018). The ARTIC network protocol was designed for nanopore technology (Oxford Nanopore Technologies), enabling rapid genome sequencing for outbreak response. The method was originally capable of testing only 23 samples plus a negative control on a flowcell, however, with the recent release of the Native Barcoding Expansion 96 kit by ONT, 11-95 samples plus a negative control can be sequenced on a flowcell using the ARTIC LoCost V3 method (Quick, 2020). A platform agnostic method is required to provide flexible throughput on Illumina or nanopore that allows low-cost sequencing of 10s to 100s of viral genomes depending on (1) changing requirements as the pandemic peaks and troughs and (2) the turnaround time required e.g. routine weekly vs rapid outbreak sequencing. Here we describe a flexible protocol, Coronavirus High Throughput (CoronaHiT), which allows for up to 95 samples, plus a negative control to be multiplexed on a single MinION flowcell or alternatively, by switching barcodes, over 384 samples on Illumina. We demonstrate CoronaHiT’s performance on 95 and 59 SARS-CoV-2 genomes on MinION and Illumina NextSeq for routine and rapid outbreak response runs, respectively, and compare to the ARTIC LoCost protocol.

## Methods

### Patient samples and RNA extraction

Samples from cases ith suspected SARS-CoV-2 were processed using five different diagnostic platforms over four laboratories in East Anglia - the Cytology Department and Microbiology Departments, NNUH, Norwich, UK, the Bob Champion Research & Education Building (BCRE), University of East Anglia, Norwich, UK and Ipswich Public Health Laboratory, Ipswich, UK.

The Cytology Department processed samples using the Roche Cobas® 8800 SARS-CoV-2 system (www.who.int/diagnostics_laboratory/eul_0504-046-00_cobas_sars_cov2_qualitative_assay_ifu.pdf?ua=1) according to the manufacturer’s instructions (n=95). The Microbiology Department processed samples using either the Hologic Panther System Aptima® SARS-CoV-2 assay (www.fda.gov/media/138096/download) (n=25) or Altona Diagnostics RealStar® SARS-CoV-s RT-PCR Kit 1.0 (altona-diagnostics.com/files/public/Content%20Homepage/-%2002%20RealStar/MAN%20-%20CE%20-%20EN/RealStar%20SARS-CoV-2%20RT-PCR%20Kit%201.0_WEB_CE_EN-S03.pdf) according to the manufacturer’s instructions (n=3). At the BCRE, RNA was extracted using the MagMAX™ Viral/Pathogen II Nucleic Acid Isolation kit (Applied Biosystems) according to the manufacturer’s instructions and the KingFisher Flex system (ThermoFisher). The presence of SARS-CoV-2 was determined using the 2019-nCoV CDC assay (https://www.fda.gov/media/134922/download) on the QuantStudio 5 (Applied Biosystems) (n=7). Ipswich Public Health Laboratory processed samples using the AusDiagnostics SARS-CoV-2, Influenza and RSV 8-well panel (www.ausdx.com/qilan/Products/20081-r01.1.pdf;jsessionid=5B2099CAE4D0D152C869A190D0032D71) (n=24). RNA was extracted from swab samples using either the AusDiagnostics MT-Prep (AusDiagnostics) or QIAsymphony (Qiagen) platforms according to the manufacturer’s instructions before being tested by the AusDiagnostics assay.

Viral transport medium from positive swabs (stored at 4°C) was collected for all samples run on the Roche Cobas®, Hologic Panther System and Altona RealStar®. In all other cases excess RNA was collected (frozen at −80°C). Excess positive SARS-CoV-2 inactivated swab samples (200µl viral transport medium from nose and throat swabs inactivated in 200 µl Zymo DNA/RNA shield and 800 µl Zymo viral DNA/RNA buffer) were collected from Cytology and the Microbiology Departments. SARS-CoV-2 positive RNA extracts (∼20 µl) were collected from Ipswich Public Health Laboratory and the BCRE as part of the COG-UK Consortium project (PHE Research Ethics and Governance Group R&D ref no NR0195). RNA was extracted from inactivated swab samples using the Quick DNA/RNA Viral Magbead kit from step 2 of the DNA/RNA purification protocol (Zymo) (files.zymoresearch.com/protocols/_r2140_r2141_quick-dna-rna_viral_magbead.pdf).

The lower of the cycle thresholds (Ct) produced by the two SARS-CoV-2 assays in the Roche, AusDiagnostics, Altona Diagnostics and CDC assays were used to determine whether samples required dilution before sequencing according to the ARTIC protocol. The Aptima SARS-CoV-2 assay on the Hologic Panther System does not provide a Ct value but rather a combined fluorescence signal for both targets in relative light units (RLUs), therefore all samples tested by the Hologic Panther were processed undiluted in the ARTIC protocol.

### ARTIC SARS-CoV-2 multiplex tiling PCR

cDNA and multiplex PCR reactions were prepared following the ARTIC nCoV-2019 sequencing protocol V3 (LoCost) (Quick, 2020). Dilutions of RNA were prepared when required based on Ct values following the guidelines from the ARTIC protocol.

V3 CoV-2 primer scheme (https://github.com/artic-network/artic-ncov2019/tree/master/primer_schemes/nCoV-2019/V3) were used to perform the multiplex PCR for SARS-CoV-2 according to the ARTIC protocol (Quick, 2020). For the ARTIC multiplex PCR, 65°C was chosen as the annealing/extension temperature, and due to variable Ct values, all samples were run for 35 cycles in the two multiplex PCRs.

### CoronaHiT-ONT library preparation

Libraries were prepared using a novel modified Illumina DNA prep tagmentation approach (formerly called Nextera DNA Flex Illumina Library Prep) (Rowan et al. 2019; Beier et al. 2017). Primers with a 3’ end compatible with the Nextera transposon insert and a 24bp barcode at the 5’ end with a 7 bp spacer were used to PCR barcode the tagmented ARTIC PCR products. The barcode sequences are from the PCR Barcoding Expansion 1-96 kit (EXP-PBC096, Oxford Nanopore Technologies). Symmetrical dual barcoding was used, i.e. the same barcode added at each end of the PCR product and up to 96 samples could be run together using this approach or 95 if a negative control is included (Supplementary Table 4).

ARTIC PCR products were diluted 1:5 (2.5 µl Pool 1, 2.5 µl Pool 2 and 20 µl PCR grade water). Tagmentation was performed as follows; 0.5 µl TB1 Tagmentation Buffer 1 , 0.5 µl BLT Bead-Linked Transposase (both contained in Illumina® DNA Prep, (M) Tagmentation Catalogue No 20018704) and 4 µl PCR grade water was made as a master mix scaled to sample number. On ice, 5 µl of tagmentation mix was added to each well of a chilled 96-well plate. Next, 2 µl of diluted PCR product was pipette mixed with the 5 µl tagmentation mix. This plate was sealed and briefly centrifuged before incubation at 55°C for 15 minutes in a thermal cycler (heated lid 65°C) and held at 10°C.

PCR barcoding was performed using Kapa 2G Robust PCR kit (Sigma Catalogue No. KK5005) as follows: 4 µl Reaction buffer (GC), 0.4 µl dNTP’s, 0.08 µl Kapa 2G Robust Polymerase and 7.52 µl PCR grade water per sample were mixed and 12 µl was added to each well in a new 96-well plate. 1 µl of the appropriate barcode pair (Supplementary Table 4) at 10µM was added to each well. Finally, the 7 µl of Tagmentation mix was added, making sure to transfer all the beads. PCR reactions were run at 72°C for 3 minutes, 95°C for 1 minute, followed by 14 cycles of 95°C for 10 seconds, 55°C for 20 seconds and 72°C for 1 minute. Following PCR, 2 µl of each sample was pooled and 40 µl of this pool was bead washed with 36 µl (0.8X) AMPure XP beads (2 washes in 200μl 70% ethanol) for the routine samples. For the rapid response run, 100 µl of the pool was washed with 60 µl (0.6X) AMPure XP. Pools were eluted in 20 µl of EB (Qiagen Catalogue No. 19086). The barcoded pool was quantified using Qubit High Sensitivity kit (Catalogue No. Q32851).

A nanopore sequencing library was then made, largely following the SQK-LSK109 protocol. The end-prep reaction was prepared as follows: 7 µl Ultra II end prep buffer, 3 µl Ultra II end prep enzyme mix, 40 µl nuclease free water and 10 µl of washed barcoded pool from the previous step (final volume 60 µl). The reaction was incubated at room temperature for 15 mins and 65°C for 10 mins, followed by a hold at 4°C for at least 1 min. This was bead-washed using 60 µl of AMPure Beads (1X) and two 200μl 70% ethanol washes and eluted in 61 µl nuclease free water. The end-prepped DNA was taken forward to the adapter ligation as follows: 30 µl end-prepped pool from previous step (∼60 ng), 30 µl nuclease free water, 25 µl LNB (ONT), 10 µl NEBNext Quick T4 Ligase and 5 µl AMX (ONT) was mixed and incubated at room temperature for 20 minutes.

After the incubation, the full volume was washed with 40 µl AMPure XP beads and 2 consecutive 250 µl SFB (ONT) washes with resuspension of beads both times and this was eluted in 15 µl of EB (ONT). The final library was quantified with Qubit High Sensitivity and size checked on a Tapestation with D5000 tape. 12 µl (∼30-50 ng) was used for flowcell loading, with the addition of 37.5 µl SQB and 25.5 µl LB.

### CoronaHiT-Illumina library preparation

PCR products were tagmented and barcoded as described for the CoronaHiT-ONT library preparation, however, standard Nextera XT Index Kit indexes were used (Sets A to D for up to 384 combinations, Illumina Catalogue No’s FC-131-2001, FC-131-2002, FC-131-2003 and FC-131-2004). The PCR master mix was adjusted and water removed to add 2 µl each of the P7 and P5 primers. Five microliters of each barcoded sample was pooled (without quantification) and 100 µl of the library pool was size selected with 0.8X AMPure XP beads (80 µl), with final elution in 50 µl EB (10mM Tris-HCl). The barcoded pool was sized on a Agilent Tapestation D5000 tape and quantified using QuantiFluor® ONE dsDNA System (Promega, WI, USA) and the molarity calculated.

The Illumina library pool was run at a final concentration of 1.5 pM on an Illumina Nextseq500 instrument using a Mid Output Flowcell (NSQ® 500 Mid Output KT v2 (300 CYS) Illumina Catalogue FC-404-2003) following the Illumina recommended denaturation and loading recommendations which included a 1% PhiX spike (PhiX Control v3 Illumina Catalogue FC-110-3001).

### ARTIC LoCost protocol Nanopore library preparation

After ARTIC multiplex PCR, library preparation was performed using the nCoV-2019 sequencing protocol v3 (LoCost) V3 (Quick, 2020). Briefly, PCR Pool 1 and 2 were pooled for each sample and diluted 1 in 10 (2.5 µl Pool 1, 2.5 µl Pool 2 and 45 µl nuclease free water), and end-prepped as follows: 1.2 µl Ultra II end prep buffer, 0.5 µl Ultra II end prep enzyme mix, 3.3 µl PCR dilution from previous step and 5 µl nuclease free water (final volume 15 µl). The reaction was incubated at room temperature for 15 min and 65°C in a thermocycler for 15 min and incubated on ice for 1 min. Native barcode ligation was prepared in a new plate: 0.75 µl end-prepped DNA, 1.25 µl native barcode, 5 µl Blunt/TA Ligase Master Mix, 3 µl nuclease free water, (final volume 10 µl). The reaction was incubated at room temperature 20 min and 65°C in a thermocycler for 10 min and incubated on ice for 1 min. Amplicons were pooled together (2 µl for 95 samples and 5 µl for 59 samples) and underwent a 0.4X AMPure bead wash with two 250 µl SFB washes and one 70% ethanol wash. DNA was eluted in 30 µl of Qiagen EB. Adapter ligation was performed on the full volume (30 µl barcoded amplicon pool, 5 µl Adapter Mix II (ONT), 10 µl NEBNext Quick Ligation Reaction Buffer (5X), 5 µl Quick T4 DNA Ligase). The ligation reaction was incubated at room temperature for 20 min and 1X bead washed (50 µl AMPure XP beads) with 250 µl SFB two times. The library was eluted in 15 µl of elution buffer (ONT) and quantified. 15 ng of the adapted library was used for final loading.

### Nanopore sequence analysis

Basecalling was performed using Guppy v.4.2.2 (Oxford Nanopore Technologies) in high accuracy mode (model dna_r9.4.1_450bps_hac), on a private OpenStack cloud at Quadram Institute Bioscience using multiple Ubuntu v18.04 virtual machines running Nvidia T4 GPU.

The CoronaHiT-ONT sequencing data were demultiplexed using guppy_barcoder (v4.2.2) with a custom arrangement of the barcodes as described at https://github.com/quadram-institute-bioscience/coronahit_guppy, with the option ‘require_barcodes_both_ends’ and a score of 60 at both ends to produce 95 FASTQ files (94 SARS-CoV-2 samples and 1 negative control) and 61 FASTQ files (59 SARS-CoV-2 samples and 2 negative control) for the routine and rapid response runs, respectively. The ARTIC ONT sequencing data were demultiplexed using guppy_barcoder (v4.2.2) with the option ‘require_barcodes_both_ends’ and a score of 60 at both ends to produce 95 FASTQ files (94 SARS-CoV-2 samples and 1 negative control) and 61 FASTQ files (59 SARS-CoV-2 samples and 2 negative control) for the routine and rapid response runs, respectively.

The downstream analysis was performed using a copy of the ARTIC pipeline (v1.1.3) as previously described (Loman, Rowe, and Rambaut 2020) to generate a consensus sequence for each sample in FASTA format. The pipeline includes the following main steps: The input reads were filtered based on reads length (ARTIC: 400-700; CoronaHiT: 150-600), and mapped to the Wuhan-Hu-1 reference genome (accession MN908947.3) using minimap2 (v 2.17-r941). The mapped bases in BAM format were trimmed off in primer regions by the ARTIC subcommand align_trim for ARTIC LoCost data. For CoronaHiT-ONT data, we used the subcommand samtools ampliconclip (v 1.11) at the primer trimming step (https://github.com/quadram-institute-bioscience/fieldbioinformatics/tree/coronahit). The trimmed reads were then used for variant calling with medaka (v 1.2.0) and longshot (v 0.4.1). The final consensus was generated from a filtered VCF file and a mask file of positions with either a depth of coverage lower than 20 or a SNP in an amplifying primer site. The consensus sequences were uploaded to GISAID and the raw sequence data was uploaded to the European Nucleotide Archive under BioProject PRJEB41737. The accession numbers for each sample are available in Supplementary Table 1. The metrics and results of all experiments are available in Supplementary Table 2 and are summarised in Table 1.

**Table 1:**
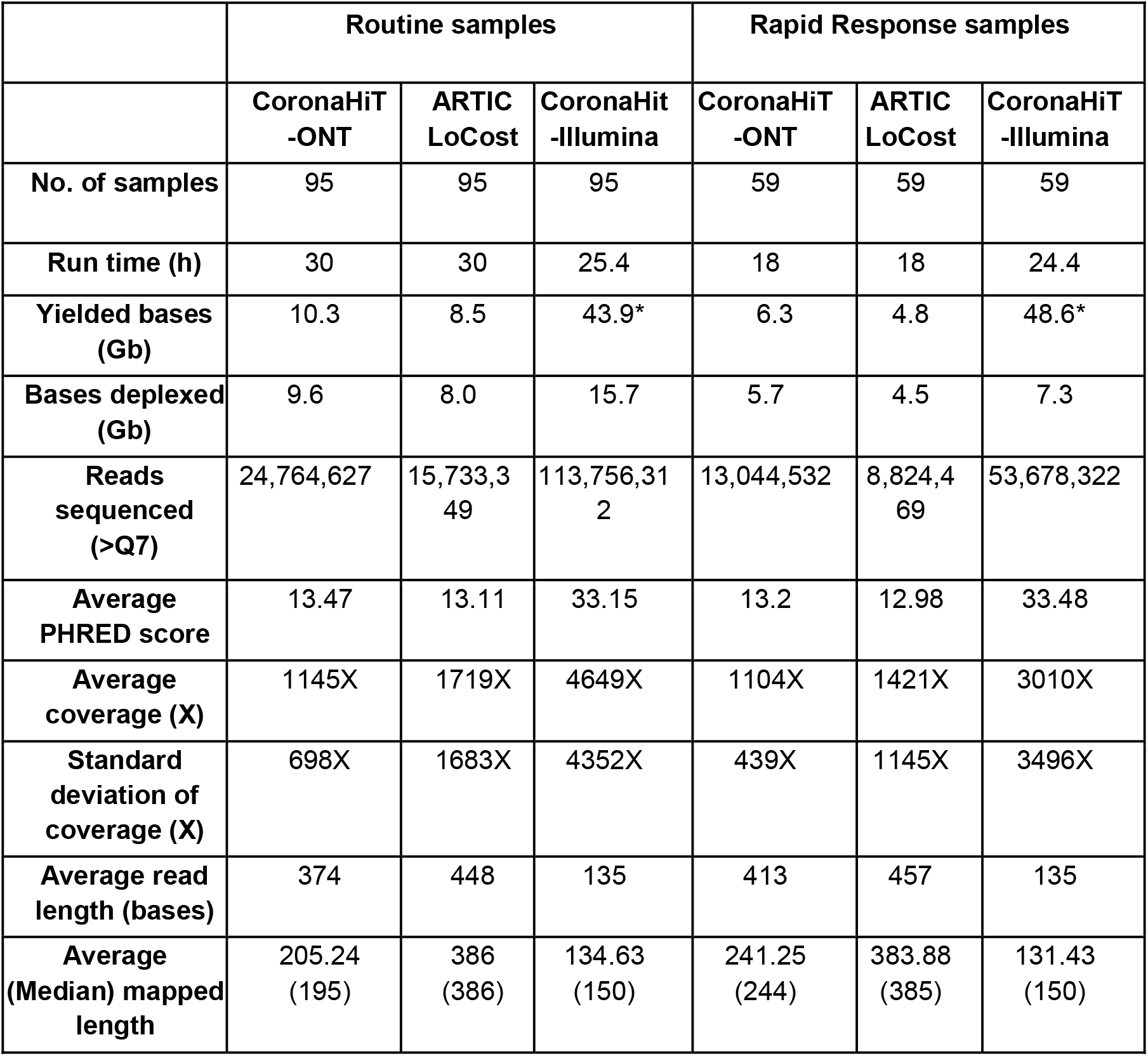
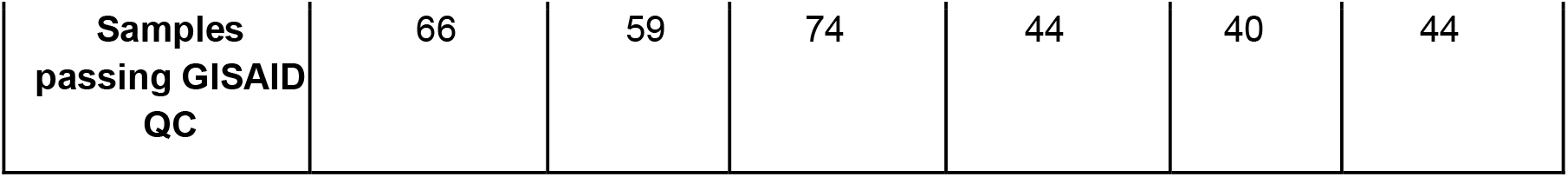
Summary statistics for each sequencing experiment. Sample specific metrics are available in Supplementary Table 2. (*The CoronaHiT-Illumina total yield includes non-relevant samples on the sequencing run, while the deplexed yield only relates to relevant samples).

|  | Routine samples |  |  | Rapid Response samples |  |  |
| --- | --- | --- | --- | --- | --- | --- |
|  | CoronaHiT-ONT | ARTIC LoCost | CoronaHiT-Illumina | CoronaHiT-ONT | ARTIC LoCost | CoronaHiT-Illumina |
| **No. of samples** | 95 | 95 | 95 | 59 | 59 | 59 |
| **Run time (h)** | 30 | 30 | 25.4 | 18 | 18 | 24.4 |
| **Yielded bases (Gb)** | 10.3 | 8.5 | 43.9* | 6.3 | 4.8 | 48.6* |
| **Bases deplexed (Gb)** | 9.6 | 8.0 | 15.7 | 5.7 | 4.5 | 7.3 |
| **Reads sequenced (>Q7)** | 24,764,627 | 15,733,349 | 113,756,312 | 13,044,532 | 8,824,469 | 53,678,322 |
| **Average PHRED score** | 13.47 | 13.11 | 33.15 | 13.2 | 12.98 | 33.48 |
| **Average coverage (X)** | 1145X | 1719X | 4649X | 1104X | 1421X | 3010X |
| **Standard deviation of coverage (X)** | 698X | 1683X | 4352X | 439X | 1145X | 3496X |
| **Average read length (bases)** | 374 | 448 | 135 | 413 | 457 | 135 |
| **Average (Median) mapped length** | 205.24 (195) | 386 (386) | 134.63 (150) | 241.25 (244) | 383.88 (385) | 131.43 (150) |
| **Samples**<br>**passing GISAID**<br>**QC** | 66 | 59 | 74 | 44 | 40 | 44 |

**Table 2:** The number of consensus genomes passing and failing the different QC thresholds for each experiment. Extended data are available in Supplementary Table 2.

|  | Routine samples |  |  | Rapid Response samples |  |  |
| --- | --- | --- | --- | --- | --- | --- |
|  | CoronaHiT-ONT | ARTIC LoCost | CoronaHiT-Illumina | CoronaHiT-ONT | ARTIC LoCost | CoronaHiT-Illumina |
| **No. of samples sequenced** | **95** | **95** | **95** | **59** | **59** | **59** |
| Consensus genomes | 98.95% (94) | 96.84% (92) | 100% (95) | 96.61% (57) | 91.53% (54) | 100% (59) |
| Passing COG-UK QC | 80.00% (76) | 76.84% (73) | 82.11% (78) | 81.36% (48) | 74.58% (44) | 81.36% (48) |
| Passing GISAID QC | 69.47% (66) | 62.11% (59) | 77.89% (74) | 74.58% (44) | 67.80% (40) | 74.58%(44) |
| Failing COG-UK QC | 20.00% (19) | 23.16% (22) | 17.89% (17) | 18.64% (11) | 25.42%(15) | 18.64% (11) |
| Failing GISAID QC | 30.53%(29) | 37.89% (36) | 22.11% (21) | 25.42%(15) | 32.20% (19) | 25.42% (15) |
| Avg (Median) Ns of COG-UK passed | 1504 (354) | 1635 (121) | 688 (53) | 977 (606) | 1101 (339) | 911 (292) |
| Avg SNPs of COG-UK passed | 7.99 | 7.99 | 11.0 | 18.3 | 18.2 | 20.4 |
| **No. of samples with known Ct $\leq 32$** | **65** | **65** | **65** | **18** | **18** | **18** |
| Consensus genomes | 100% (65) | 100% (65) | 100% (65) | 100% (18) | 100% (18) | 100% (18) |
| Passing COG-UK QC | 98.46% (64) | 98.46% (64) | 98.46% (64) | 100%(18) | 94.44% (17) | 100% (18) |
| Passing GISAID QC | 95.38% (62) | 89.23% (58) | 98.46% (64) | 94.44% (17) | 88.89% (16) | 94.44% (17) |
| Failing COG-UK QC | 1.54% (1) | 1.54% (1) | 1.54% (1) | 0% (0) | 5.56% (1) | 0% (0) |
| Failing GISAID QC | 4.62% (3) | 10.77% (7) | 1.54% (1) | 5.56% (1) | 11.11% (2) | 5.56% (1) |
| Avg (Median) Ns of COG-UK passed | 682 (339) | 815 (121) | 111 (47) | 895 (339) | 911 (121) | 1064 (514) |
| Avg SNPs of COG-UK passed | 8.19 | 8.17 | 10.2 | 18.8 | 18.9 | 20 |

### Illumina sequence analysis

Additional samples, not reported in this study, were included on Illumina NextSeq runs. The raw reads were demultiplexed using bcl2fastq (v2.20) (Illumina Inc.) to produce 311 FASTQ files for the run with the routine samples (112 SARS-CoV-2 samples and 3 negative controls) and the run with the rapid response samples (247 SARS-CoV-2 samples, 4 negative controls, and 2 positive controls) with only the relevant samples analysed in this paper. The reads were used to generate a consensus sequence for each sample using an open source pipeline adapted from https://github.com/connorlab/ncov2019-artic-nf (https://github.com/quadram-institute-bioscience/ncov2019-artic-nf/tree/qib). Briefly, the reads had adapters trimmed with TrimGalore (https://github.com/FelixKrueger/TrimGalore), were aligned to the Wuhan-Hu-1 reference genome (accession MN908947.3) using BWA-MEM (v0.7.17) (Li 2013), the ARTIC amplicons were trimmed and a consensus built using iVAR (v.1.2.3) (Grubaugh et al. 2019).

### Quality Control

The COG-UK consortium defined a consensus sequence as passing COG-UK quality control if greater than 50% of the genome was covered by confident calls or there was at least 1 contiguous sequence of more than 10,000 bases and with no evidence of contamination. This is regarded as the minimum amount of data to be phylogenetically useful. A confident call was defined as having a minimum of 10X depth of coverage for Illumina data and 20X depth of coverage for Nanopore data. If the coverage fell below these thresholds, the bases were masked with Ns. Low quality variants were also masked with Ns. The QC threshold for inclusion in GISAID was higher, requiring that greater than 90% of the genome was covered by confident calls with no evidence of contamination.

### Phylogenetic analysis

For each sample sequenced in 3 separate experiments (CoronaHiT-ONT, CoronaHiT-Illumina, ARTIC-ONT), a phylogeny was generated from all of the consensus genomes (n=216 for the routine samples and n=132 for the rapid response samples) passing GISAID QC over all experiments (n=72 out of 95, and n=44 out of 59). A multiple FASTA alignment was created by aligning all samples to the reference genome MN908947.3 with MAFFT v7.470. A maximum likelihood tree was estimated with IQTREE2 (v2.0.4) (Minh et al. 2020) under the HKY model (Hasegawa, Kishino, and Yano 1985), collapsing branches smaller than 10^−7^ into a polytomy. SNPs in the multiple FASTA alignment were identified using SNP-sites (v2.5.1) (Page et al. 2016) and the tree was visualised with FigTree (v1.4.4) (https://github.com/rambaut/figtree).

## Results

A novel library preparation method, CoronaHiT, was developed for SARS-CoV-2 genome sequencing, which combines a cheap transposase-based introduction of adapters (Illumina Nextera) with symmetric PCR barcoding of up to 96 samples (or 95 samples with a negative control) on a MinION. Alternatively, if higher throughput is needed, the barcodes can be switched for Illumina sequencing. For ONT sequencing, Nextera adapter complementary primer sequences were added to ONT PCR barcodes and used to barcode ARTIC PCR products (Figure 1) as described in the methods. For Illumina sequencing, the method is a streamlined and cheaper version of standard Illumina library preparations. CoronaHiT does not require individual sample washes and allows samples to be processed uniformly without quantification or normalisation as with the ARTIC LoCost method.

**Figure 1:**
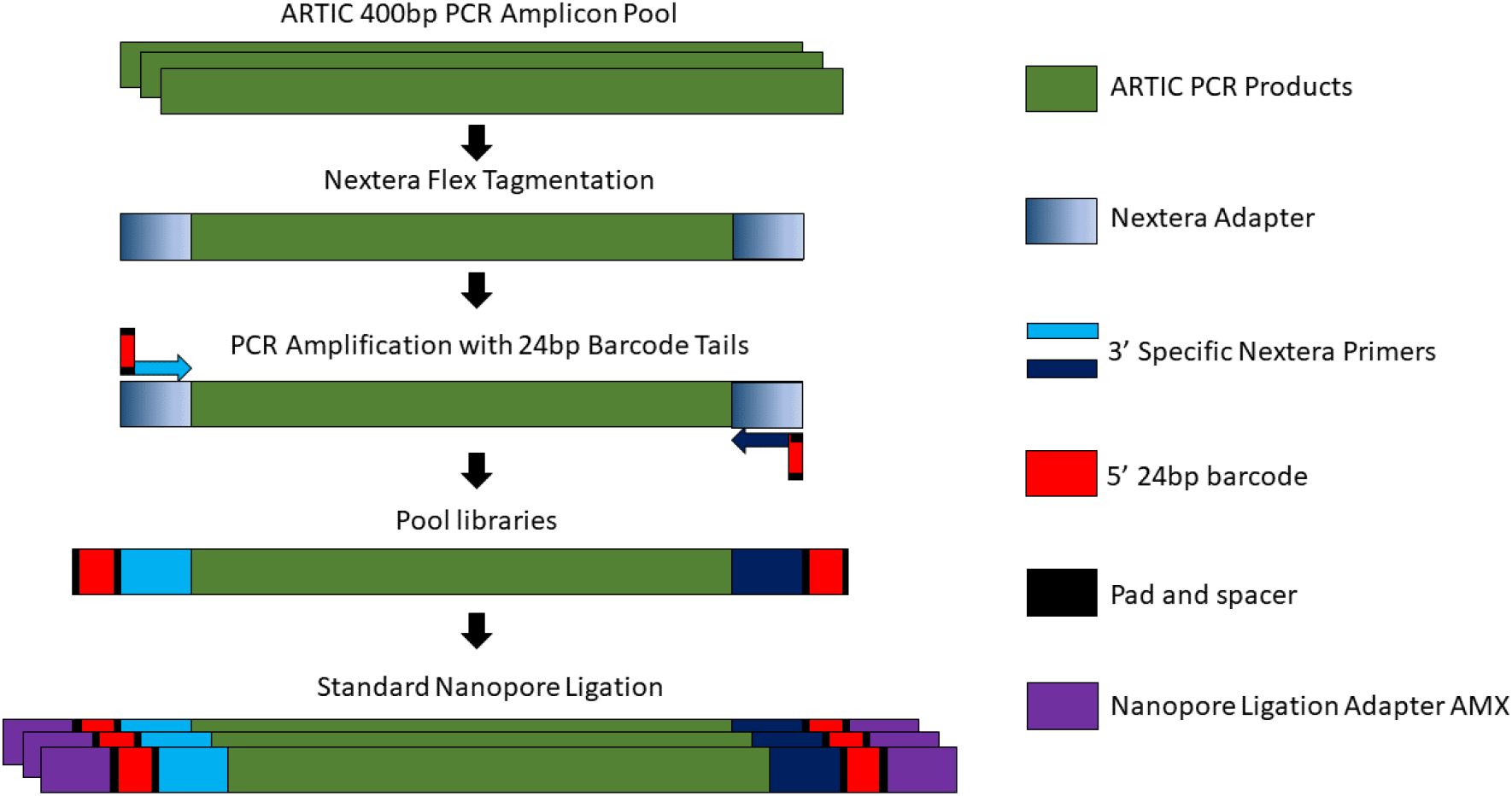
Workflow of CoronaHiT-ONT library preparation.

The CoronaHiT method was tested by multiplexing 95 SARS-CoV-2 routine COG-UK samples plus a blank (hereinafter referred to as the Routine Samples) on a MinION flowcell and on an Illumina NextSeq run. Another 59 samples, including 18 query outbreak samples, plus blanks (hereinafter referred to as the Rapid Response samples) were rapidly sequenced (within 24 hrs of receipt, with results available the following day) on a second flowcell, as well as on Illumina NextSeq. All samples were also sequenced using the ARTIC LoCost library preparation protocol on the MinION for comparison.

For the routine samples, 30 hours of sequencing data was used for both CoronaHiT-ONT and ARTIC LoCost, and for the rapid response set, 18 hours was used; the full dataset was used for both CoronaHiT-Illumina runs. The different methods produced different amounts of demultiplexed data. For the routine samples, CoronaHiT-ONT yielded 9.6 Gbases of sequence data, ARTIC LoCost sequencing produced 8.0 Gbases of data, and CoronaHiT-Illumina yielded 15.7 Gbases giving on average 1145X, 1719X and 4649X coverage per sample (Table 1). For the rapid response dataset, CoronaHiT-ONT produced 5.7 Gbases, ARTIC LoCost 4.5 Gbases, and CoronaHiT-Illumina 7.3 Gbases resulting in 1104X, 1421X, and 3010X coverage per sample respectively. Both CoronaHiT-ONT runs had less variation in coverage between samples compared to the ARTIC LoCost runs, with lower standard deviation relative to the mean (Table 1). The lower coverage for CoronaHiT-ONT compared to ARTIC is related to the shorter read lengths and the increased proportion of barcode/adapter sequence in each read and, hence, the reduced mappable region of each read.

Taking all the genomes which passed COG-UK QC, the CoronaHiT-Illumina sequencing runs produced the shortest mappable mean read length at 135 and 131 bases for the routine samples and rapid response samples respectively, just short of the maximum 150 bases for the PE 151 chemistry; ARTIC LoCost produced 386 and 384 bases, and CoronaHiT-ONT sequencing produced mappable mean read lengths of 205 and 241 bases. The shorter read lengths for CoronaHiT are related to the use of bead-linked transposases for tagmentation, resulting in the removal of the ends of the ARTIC PCR products. The introduction of a 0.6X bead wash for the rapid response CoronaHiT-ONT run (instead of the 0.8X bead wash for the routine run) resulted in the longer mapped reads and contributed to a reduction in the difference in average coverage between CoronaHiT and ARTIC (from 1145x vs 1719x in routine run dropping to 1104X vs 1421X in the rapid response run, with similar ratios of raw data produced by the methods in the two runs).

The demultiplexing steps for CoronaHiT-ONT were different from those used for ARTIC ONT sequencing as described in the methods section. Comparing the nanopore sequencing methods for the routine samples, 74.7% and 81.9% of reads were demultiplexed successfully for CoronaHiT-ONT and ARTIC LoCost respectively when only reads with a PHRED (quality) score above Q7 are considered; for the rapid response set, 69.6% and 71.6% were demultiplexed for CoronaHiT-ONT and ARTIC LoCost. The rest of the reads were unassigned, due to an inability to detect the barcode sequences at both ends of the reads. The negative controls contained zero mapping reads to SARS-CoV-2 for all nanopore datasets. The Illumina routine dataset had mapped reads, however, the vast majority were primers dimers (range of 0-4 SARS-CoV-2 reads >40bp mapped out of the 3 negative controls).

Poor quality consensus genomes were generally associated with a lower SARS-CoV-2 viral load in the clinical samples i.e. higher RT-qPCR Ct values (generally above Ct 32) were more likely to fail COG-UK and GISAID quality control thresholds. For all methods the number of Ns increased significantly in samples with a Ct above 32, which equates to approx 100 viral genome copies in the PCR reaction (Figure 2).

**Figure 2:**
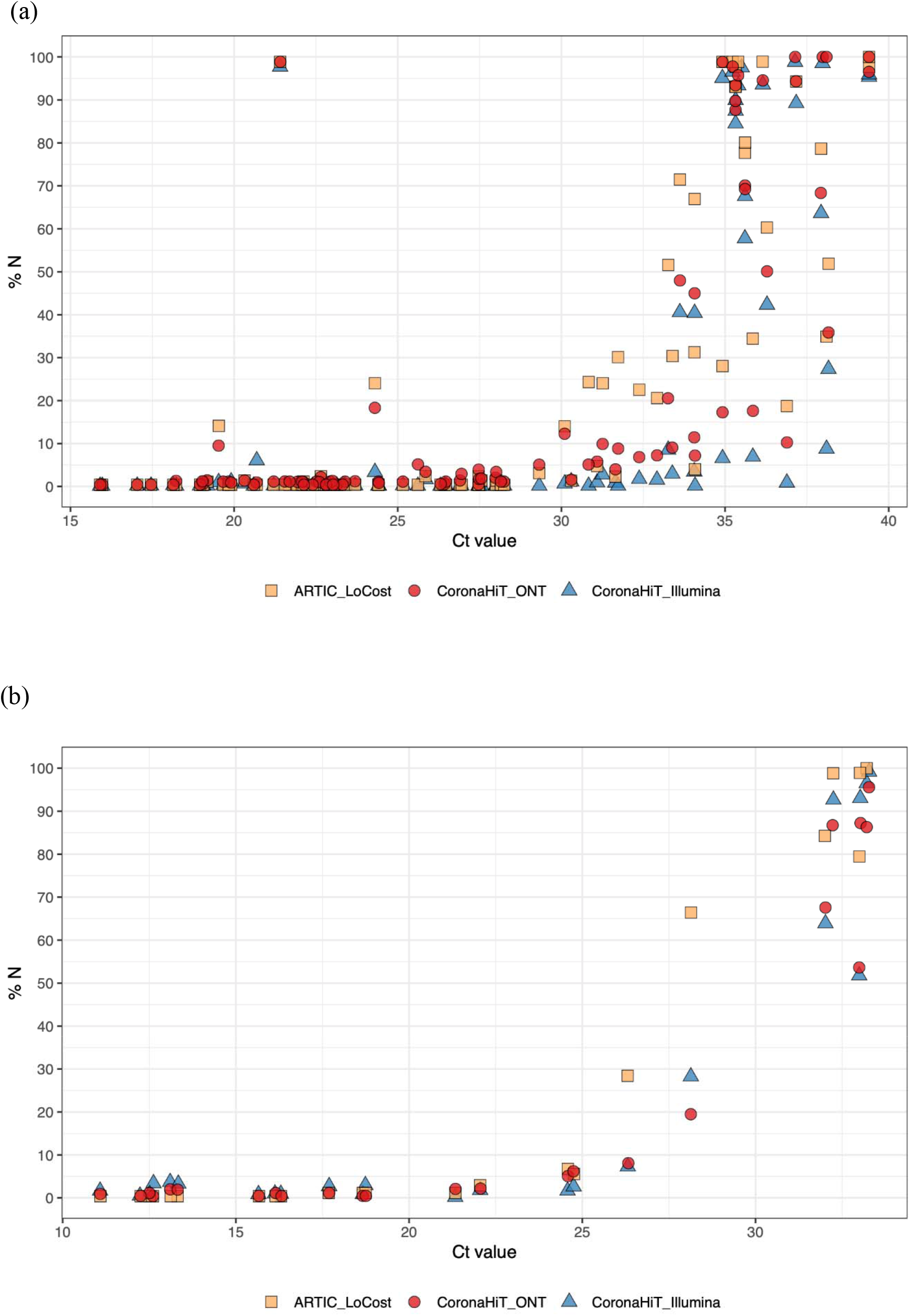
Ct value of the SARS-CoV-2 positive RNA samples sequenced using all three sequencing methods vs total number of Ns in the consensus sequence for the (a) routine sample set (b) and the rapid response sample set.

Supplementary Figures 1a-f show the Ns (missing or masked bases) within the consensus genomes - the three ARTIC PCR primer dropout areas (Benjamin Farr et al. 2020) are clearly visible. Comparing the routine samples with a Ct of 32 or below (n=65; Cts for most rapid response samples were unknown), the mean (median) number of Ns was 815 (121) for ARTIC LoCost, 111 (47) for CoronaHiT-Illumina, and 682 (339) for CoronaHiT-ONT. If all samples are included for the routine set (including higher Ct samples) then the number of Ns increases substantially to a mean (median) of 1635 (121) bases for ARTIC LoCost, 688 (53) for CoronaHiT-Illumina and 1504 (359) for CoronaHiT-ONT.

The number of samples passing the COG-UK QC criteria was 73 for ARTIC LoCost, 76 for CoronaHiT-ONT and 78 for CoronaHiT-Illumina in the routine set and 44 for ARTIC LoCost, and 48 for both CoronaHiT-ONT and CoronaHiT-Illumina in the rapid response set. The stricter GISAID QC criteria reduces the number of samples passing QC, with the CoronaHiT method outperforming ARTIC LoCost. For the routine samples, 59 samples passed for ARTIC LoCost, 66 passed for CoronaHiT-ONT and 74 passed for CoronaHiT-Illumina and for the rapid response set 40 passed for ARTIC LoCost, and 44 passed for both CoronaHiT-ONT and CoronaHiT-Illumina. Overall, the pass rate was 64.3% for ARTIC LoCost, 71.4% for CoronaHiT-ONT and 76.6% for CoronaHiT-Illumina. When considering higher viral load samples with a known Ct of 32 or below, the pass rate for both GISAID and COG-UK QC was higher, with 89.2% passing for ARTIC LoCost and 95.2% and 97.6% passing for CoronaHiT-ONT and CoronaHiT-Illumina, respectively (full details are shown in Table 2). CoronaHiT-ONT had a higher pass rate compared to ARTIC LoCost even though the average coverage was lower, this related to more even coverage across samples on the flowcell (lower standard deviation between samples relative to the mean - Table 1).

To assess the impact of data quality differences on clustering of lineages, we built maximum likelihood trees for both the routine and rapid response runs with each of the 72 and 44 consensus genomes that passed QC from the ARTIC LoCost, CoronaHiT-ONT and CoronaHiT-Illumina sequencing experiments. When the consensus genomes were placed on a phylogenetic tree for the routine set, CoronaHiT-Illumina, ARTIC LoCost, CoronaHiT-ONT showed the same clustering for most samples, except for three cases (EB1DB, EC741 and EC644) where we note that their ARTIC LoCost consensus show an increased number of ambiguous bases. All variant differences between the samples are noted in Supplementary Table 3, together with the sequence length (discounting ambiguous bases whenever there is a difference). Out of all samples in both datasets, there were only two SNP discrepancies, one in sample F04F8 between CoronaHiT-ONT and CoronaHiT-Illumina, with ARTIC LoCost calling the SNP ambiguous, and in sample F0A23 with CoronaHiT-ONT disagreeing with the other methods (Supplementary Table 3). The SNP differences did not affect the classification (i.e. closest sequence in the database), and there were no SNP differences between ARTIC-ONT and CoronaHiT-Illumina. The main other source of variation between the samples is that the Illumina genomes allow IUPAC (IUPAC-IUB Comm. on Biochem. Nomenclature (CBN) 1970) symbols for “partially” ambiguous bases. These data show that CoronaHiT provides highly accurate lineage calling compared to ARTIC LoCost.

The average number of SNPs between the Wuhan-Hu-1 reference genome and the consensus genomes varied between 7.99 SNPs for and 11.00 SNPs for the routine samples, and 18.2 and 20.4 SNPs for the rapid response samples across all methods (see Table 2 and Supplementary Table 2). The mean number of SNPs in CoronaHiT-Illumina was higher compared to the two ONT sequencing methods (Table 2) due to ambiguous bases in the Illumina dataset being regarded as SNPs in these calculations (Table 2).

The reagent cost per sample for CoronaHiT-ONT was £8.46 when sequencing 95 samples and a negative control on a MinION flowcell, marginally cheaper but similar to ARTIC sequencing at £9.75 per sample (cost breakdown in Supplementary Table 5). If 384 samples are sequenced on an Illumina NextSeq Mid output run with the CoronaHiT library preparation method, the per sample cost is £5.62.

## Discussion

Rapid viral genome sequencing during outbreaks is changing how we study disease epidemiology (Kafetzopoulou et al. 2019; Joshua Quick et al. 2016). The recent SARS-CoV-2 global pandemic has again highlighted the use of sequencing in the control of the spread of the disease. Nanopore technology is particularly suited to outbreak sequencing as it is portable, does not require expensive machinery and is accessible throughout the world (Faria et al. 2016). We present a novel platform agnostic method, CoronaHiT, for flexible throughput, cost effective and low complexity sequencing of SARS-CoV-2 genomes to respond to the pandemic at the local and national level.

The ARTIC LoCost protocol (Quick, 2020) has been widely adopted for SARS-CoV-2 genome sequencing and allows up to 95 samples (plus a negative control) to be sequenced at a time on a MinION. CoronaHiT is just as cheap, simple and fast, but the combination of transposase introduction of adapters with PCR based barcoding allows for more even coverage between multiplexed samples, resulting in a higher proportion of samples passing QC. It is also designed to be platform agnostic, simply switching barcodes to move to Illumina. This allows the user to flexibly sequence low or high throughput depending on rapidly changing requirements in the pandemic (Bayliss et al. 2017; Josh Quick 2020). With the use of asymmetric barcode primers described in Perez-

Sepulveda et al. 2020, it is possible to sequence SARS-CoV-2 at very high throughput on Illumina; in fact we have recently sequenced over 1000 SARS-CoV-2 genomes on a single Illumina NextSeq High Output run using this approach (data not shown). The CoronaHiT-Illumina library preparation method is cheaper (reduced reaction volumes) and significantly more streamlined (no sample washing or quantification before pooling, no use of stop solution, no clean-up after tagmentation and no clean-up of barcoded PCR products) than standard Illumina library preparation.

Tiling PCR approaches, such as ARTIC, are prone to high genome coverage variation due to variable primer efficiency in multiplex reactions. Some regions of the SARS-CoV-2 genome have hundreds of times higher coverage than adjacent regions using ARTIC, therefore average coverage of at least 1000X is required to obtain at least 20X coverage of the difficult regions of the genome. We demonstrate that we can achieve >1000X SARS-CoV-2 genome coverage in ∼20 minutes per sample using CoronaHiT-ONT on MinION, with a full set of 95 samples taking ∼30 hours). While the CoronaHiT-ONT runs described here are very consistent, sequencing yield depends on flowcell quality. We recommend aiming for at least 100 Mbases of estimated sequencing yield per sample to provide sufficient data for >1000X coverage/sample (average across flowcell) using CoronaHiT-ONT.

Results demonstrate that all methods are unreliable at producing high quality consensus genomes from positive clinical samples with diagnostic RT-qPCR Cts above 32 (approx. 100 viral genome copies), however, CoronaHiT performs better in these samples (Figure 2), producing fewer Ns, likely due to the additional rounds of PCR during barcoding. Below or equal to Ct 32, CoronaHiT-ONT, CoronaHiT-Illumina and ARTIC LoCost produce similar results. While more samples pass both QC measures with CoronaHiT-ONT and CoronaHiT-Illumina compared to ARTIC LoCost, primer dropout regions can be more pronounced in these methods (Supplementary Figure 1). For higher quality consensus genomes, sequencing may be run for longer. Additionally, a reduction in ARTIC PCR annealing temperature from 65°C to 63°C may help improve coverage across these regions (Benjamin Farr et al. 2020). However, data produced from CoronaHiT was sufficient to provide accurate consensus genomes that result in the same lineages and on the same branches on the phylogenetic tree as ARTIC LoCost (Figure 3). Therefore, we have demonstrated high quality, multiplexed SARS-CoV-2 genome sequencing of 95 samples on a single flowcell. If the ARTIC PCR step is optimised to even the coverage of the amplicons (as demonstrated in the Sanger COVID-19 ARTIC Illumina protocol (Benjamin Farr et al. 2020)), less overall coverage will be required per genome and more samples can be multiplexed using all methods.

**Figure 3:**
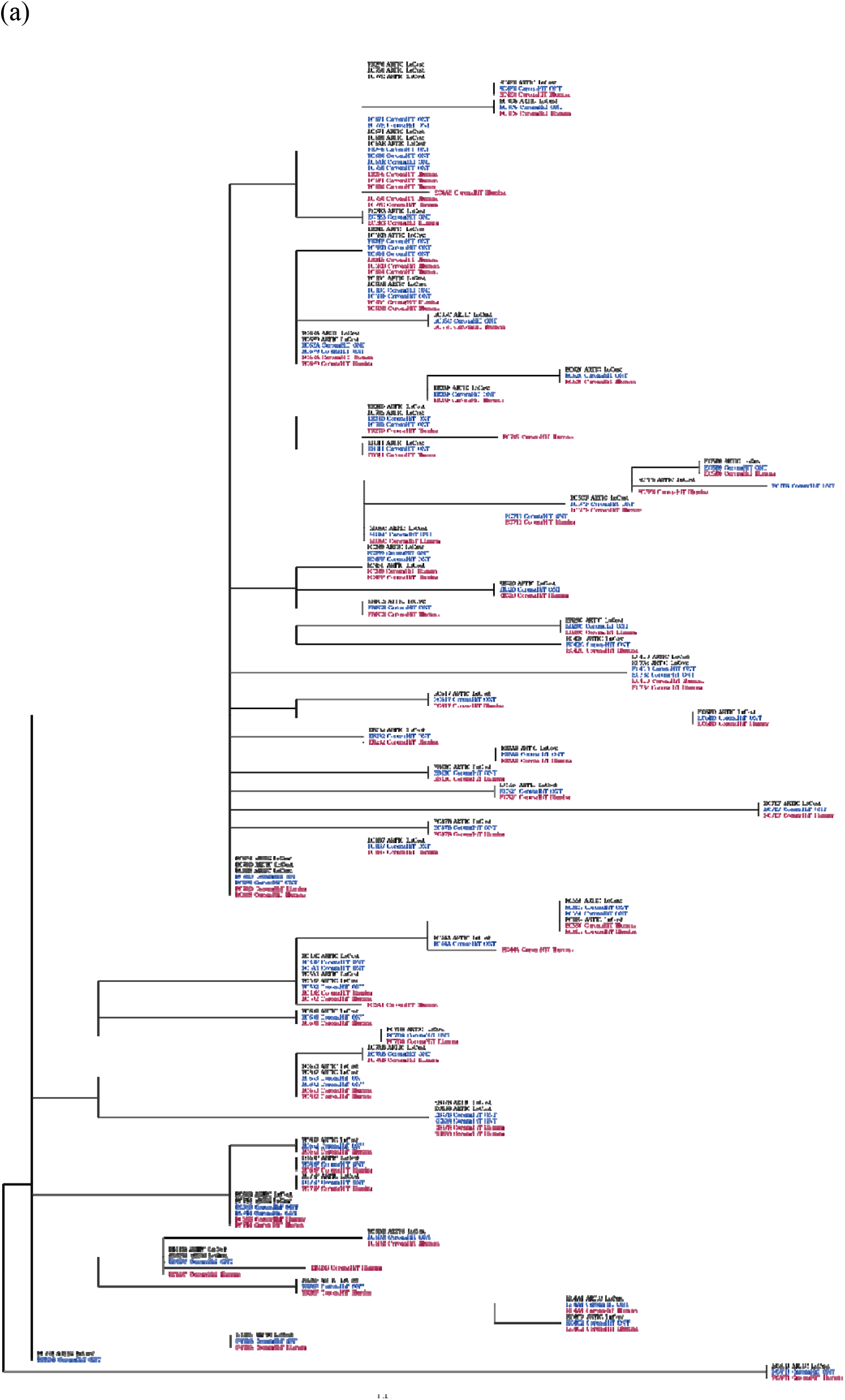
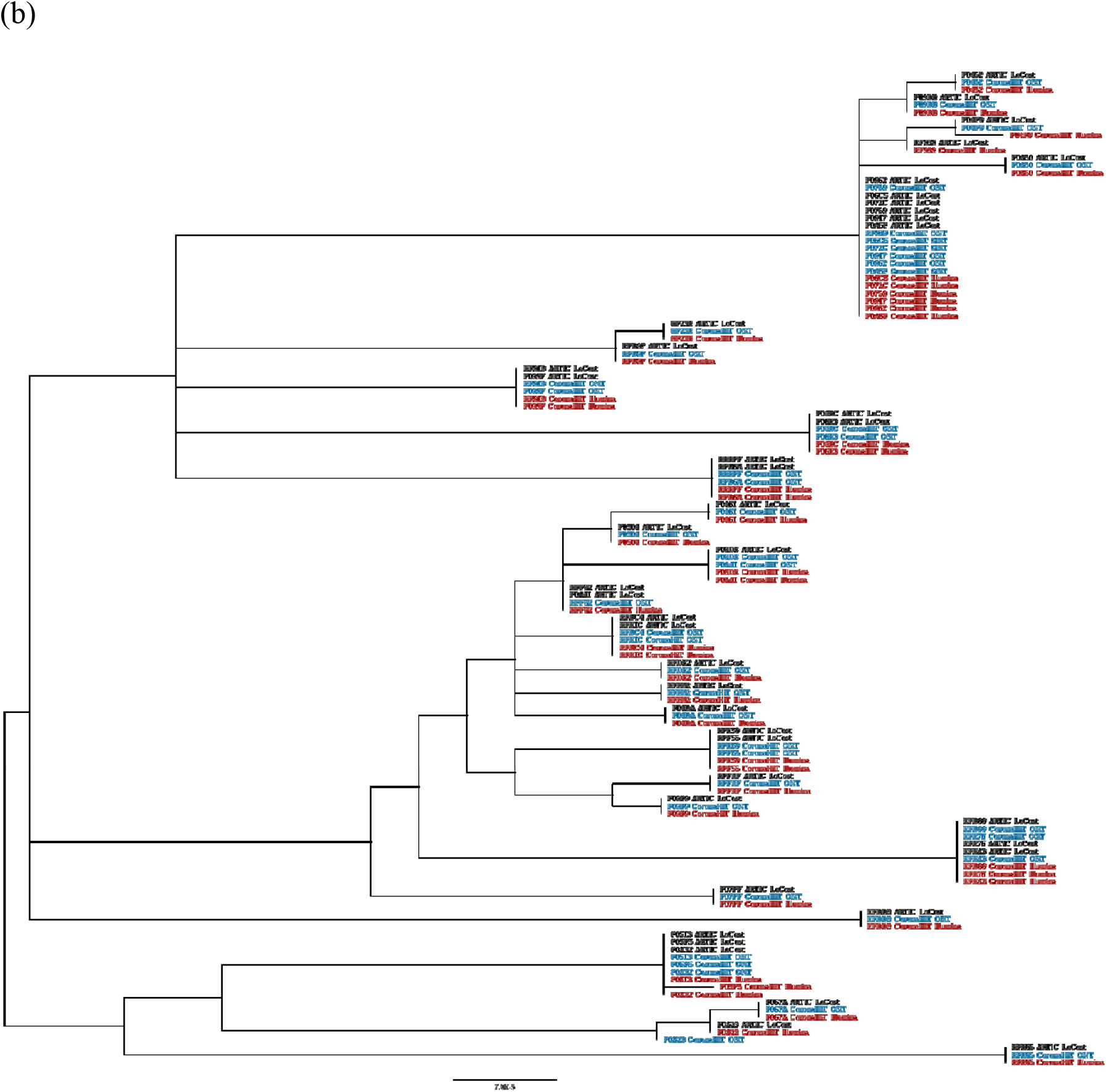
Maximum likelihood tree of the consensus genomes from each sequencing methods, showing agreement between methods for the (a) routine samples and (b) rapid response samples.

In conclusion, we demonstrate that CoronaHiT can be used to sequence 96 SARS-CoV-2 samples on a single MinION flowcell, with the option of higher throughput on Illumina. This platform agnostic method is simple, rapid and cheap and results in more samples passing QC than ARTIC LoCost while providing almost identical phylogenetic results. CoronaHiT can help scientists around the world sequence SARS-CoV-2 genomes with highly flexible throughput, thereby increasing our understanding, and reducing the spread, of the pandemic.

## Ethical approval

The COVID-19 Genomics UK Consortium has been given approval by Public Health Englands Research Ethics and Governance Group (PHE R&D Ref: NR0195).

## Acknowledgements

Thanks to George Taiaroa and Torsten Seemann from the Microbiological Diagnostic Unit Public Health Laboratory at the University of Melbourne for their advice and assistance, to Niamh Tumelty from the University of Cambridge for assistance and to Darren Heavens from the Earlham Institute for his advice on library preparation. Thanks to the COG-UK Consortium Study Group for their contributions.

## Financial declaration

LG received a partial support for his PhD from Roche. The use of Roche technology for diagnostics in NNUH is coincidental.

## Author contributions

All authors have read this manuscript and consented to its publication. The CoronaHiT method was developed by DJB and AA. The study was designed and conceived by DJB, JOG, AJP. Paper writing was by DJB, AA, AJP, JOG, GLK, APT, TLV, SR, LM.

Sequencing and library preparation was performed by DJB, AA, SR, GLK, APT, AB, AJT, NMT, RG, JOG. Bioinformatics analysis and informatics were performed by TLV, LOM, NFA, AJP. Clinical diagnostics and extractions were managed by AK, SD, RP, NE, EM. Samples and metadata were collected by LG, AB, AVG, EMA, AK and MD and biobanked by RS, RNA was extracted by AB, AJT. Risk assessments were by GLK, JW. Project management and oversight was by GLK, JOG, AJP, LM, MW, AEM, JW. Funding for the project was secured by JOG, AJP, IGC.

