## Supplementary Figures for "CoronaHiT: High throughput sequencing of SARS-CoV-2 genomes"

(a)

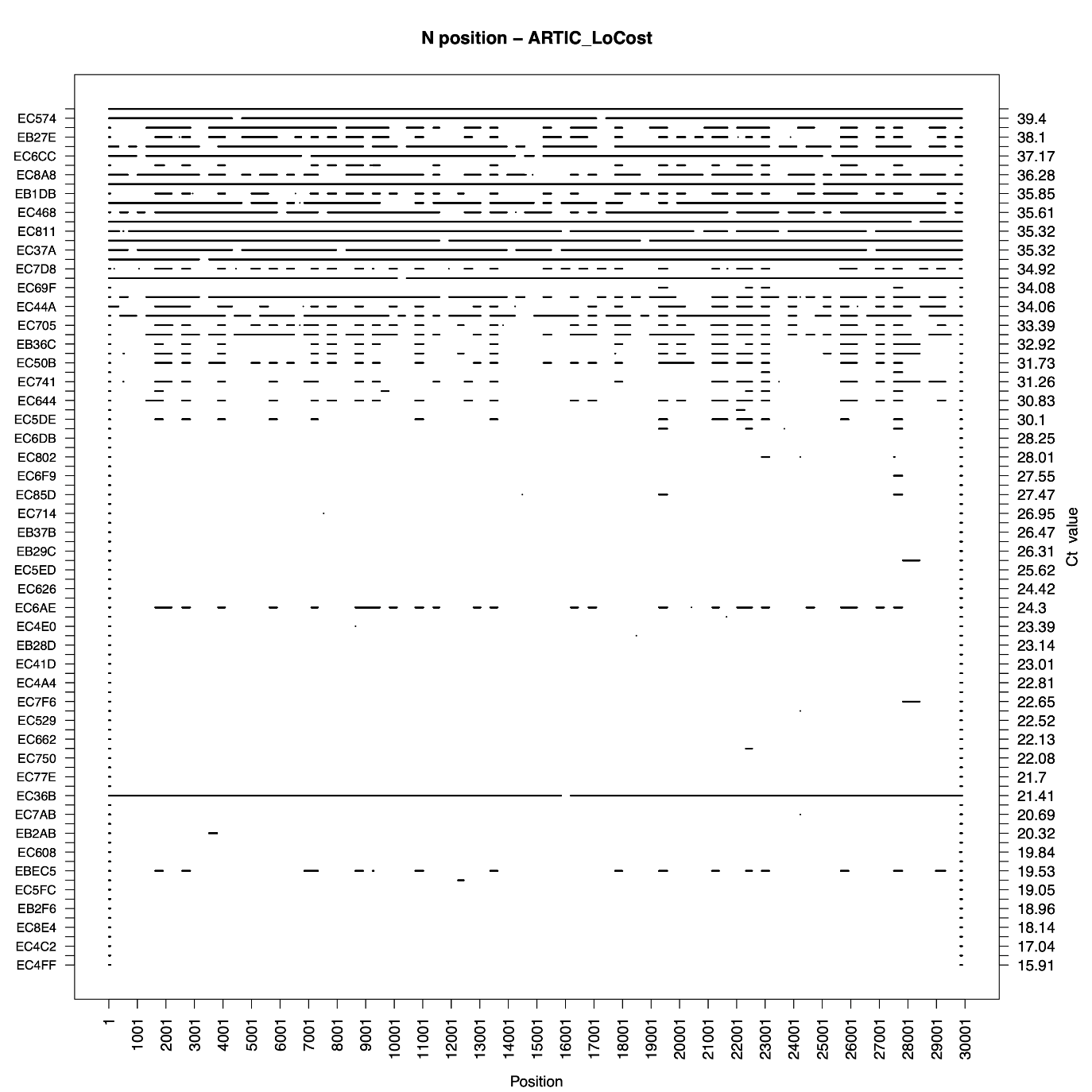

(b)

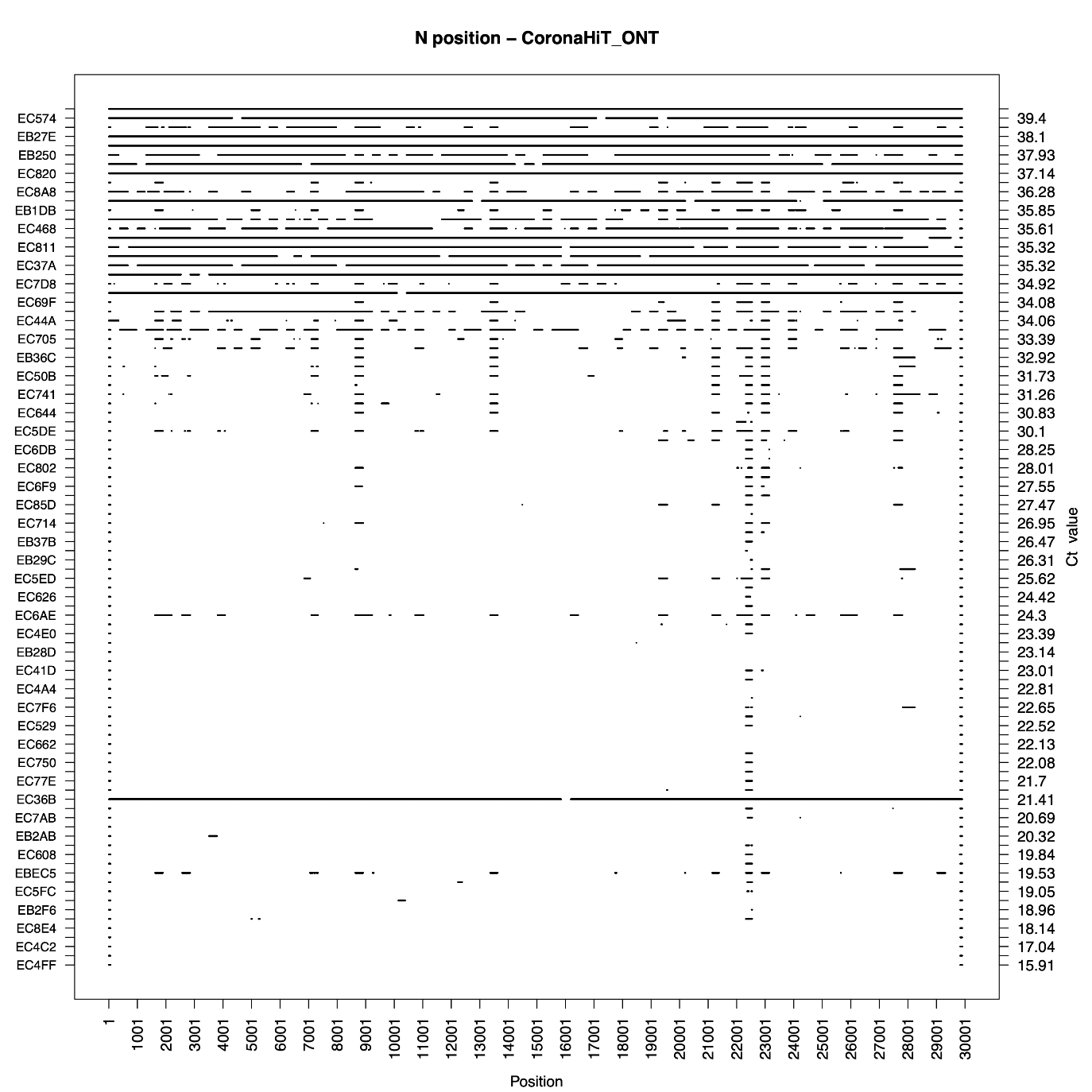

(c)

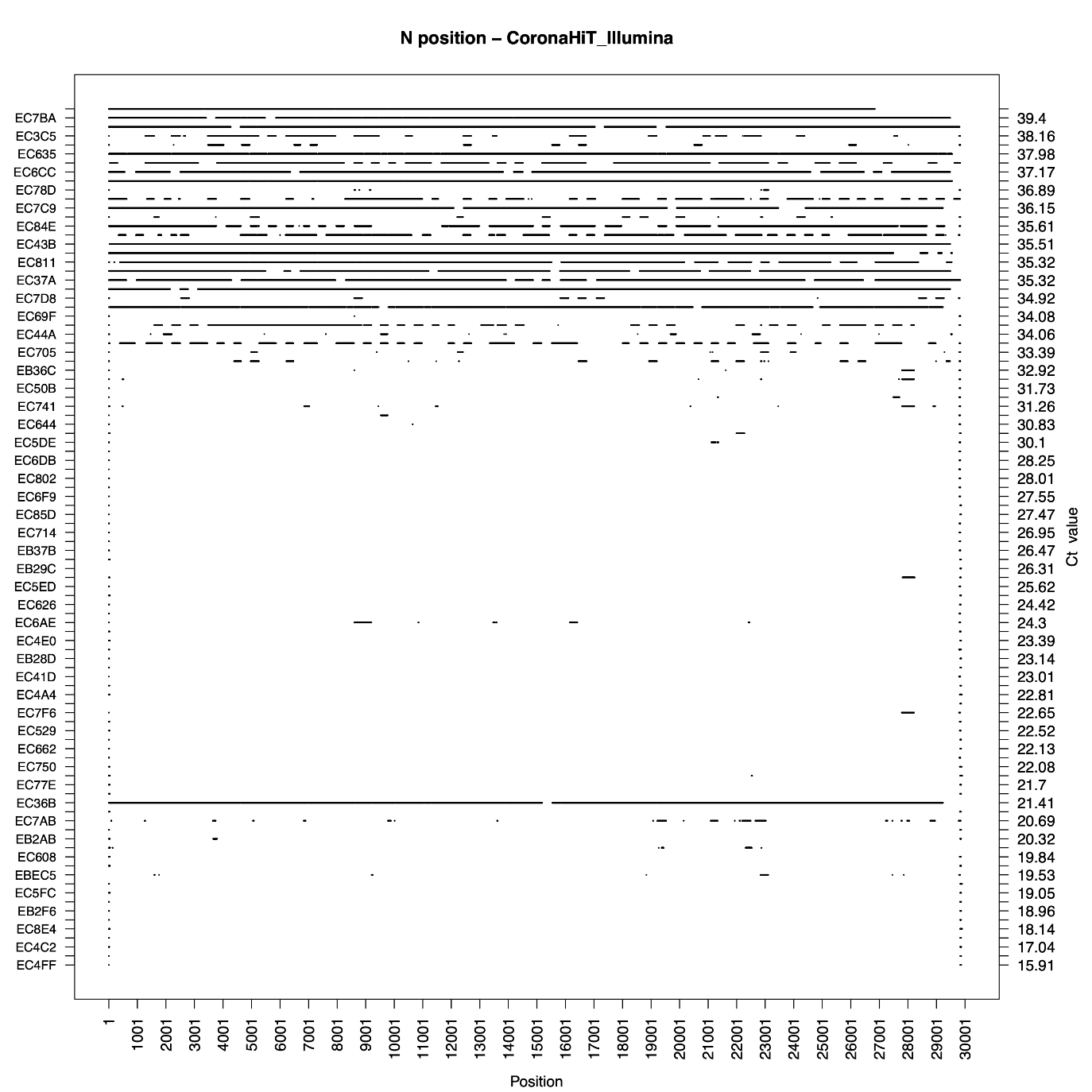

(d)

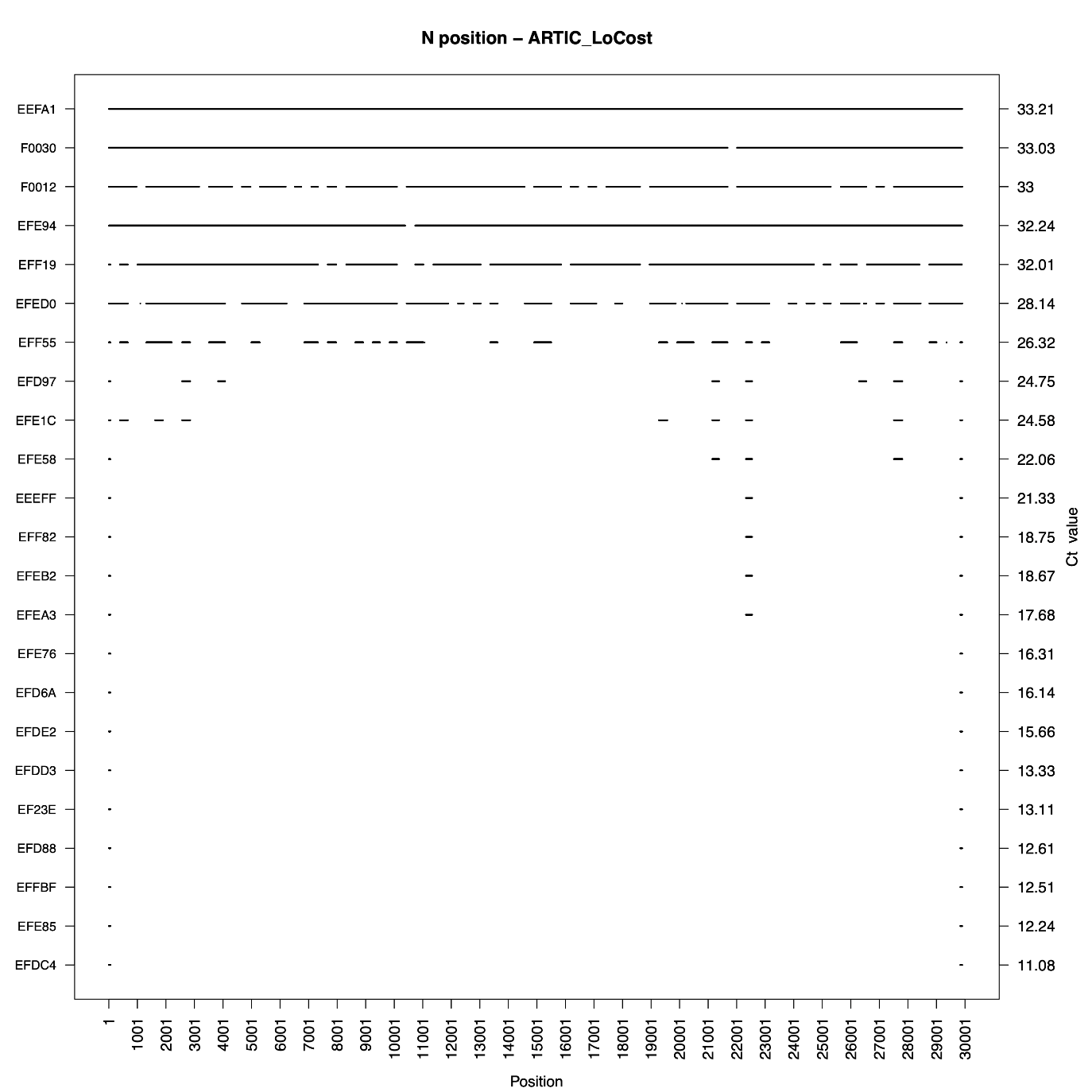

(e)

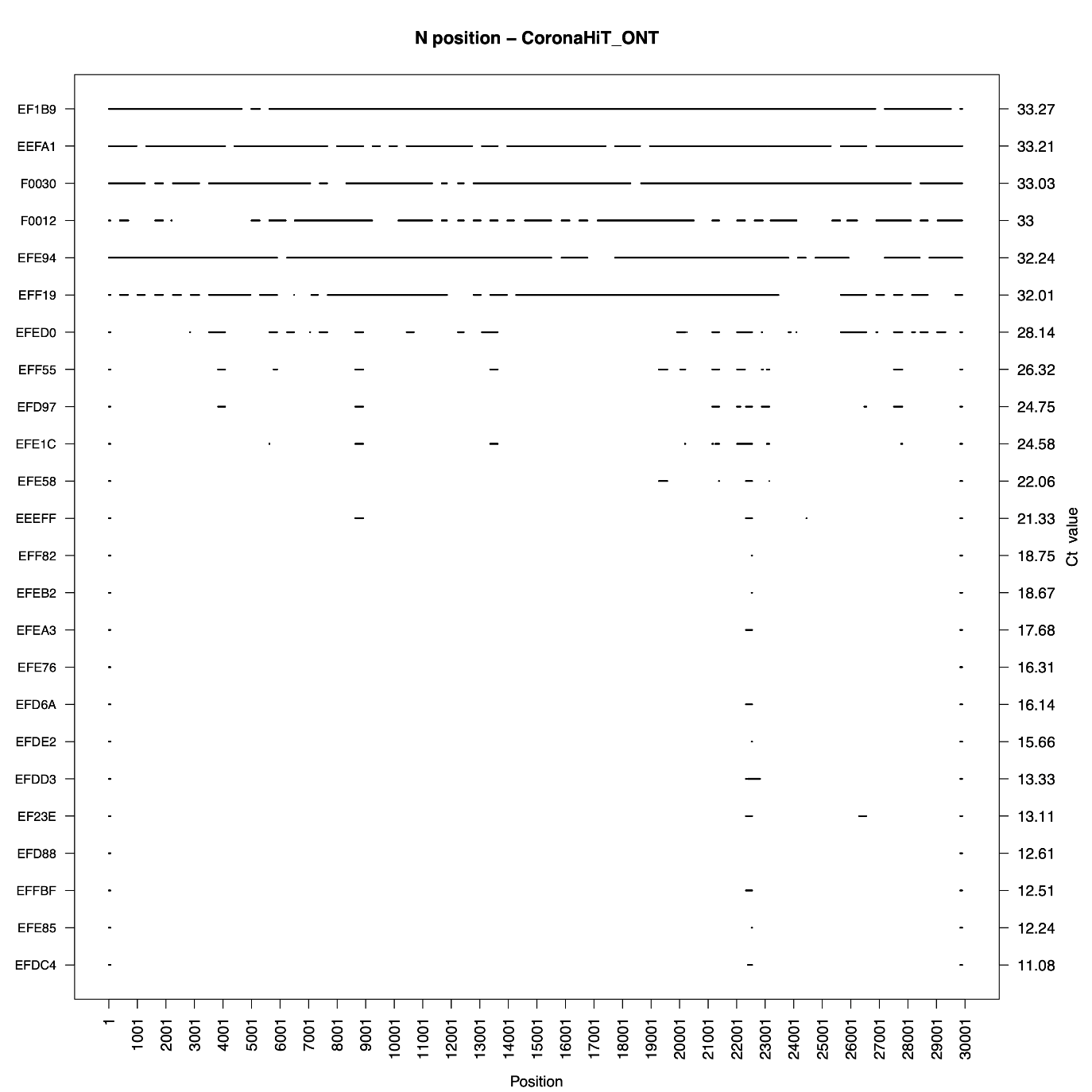

(f)

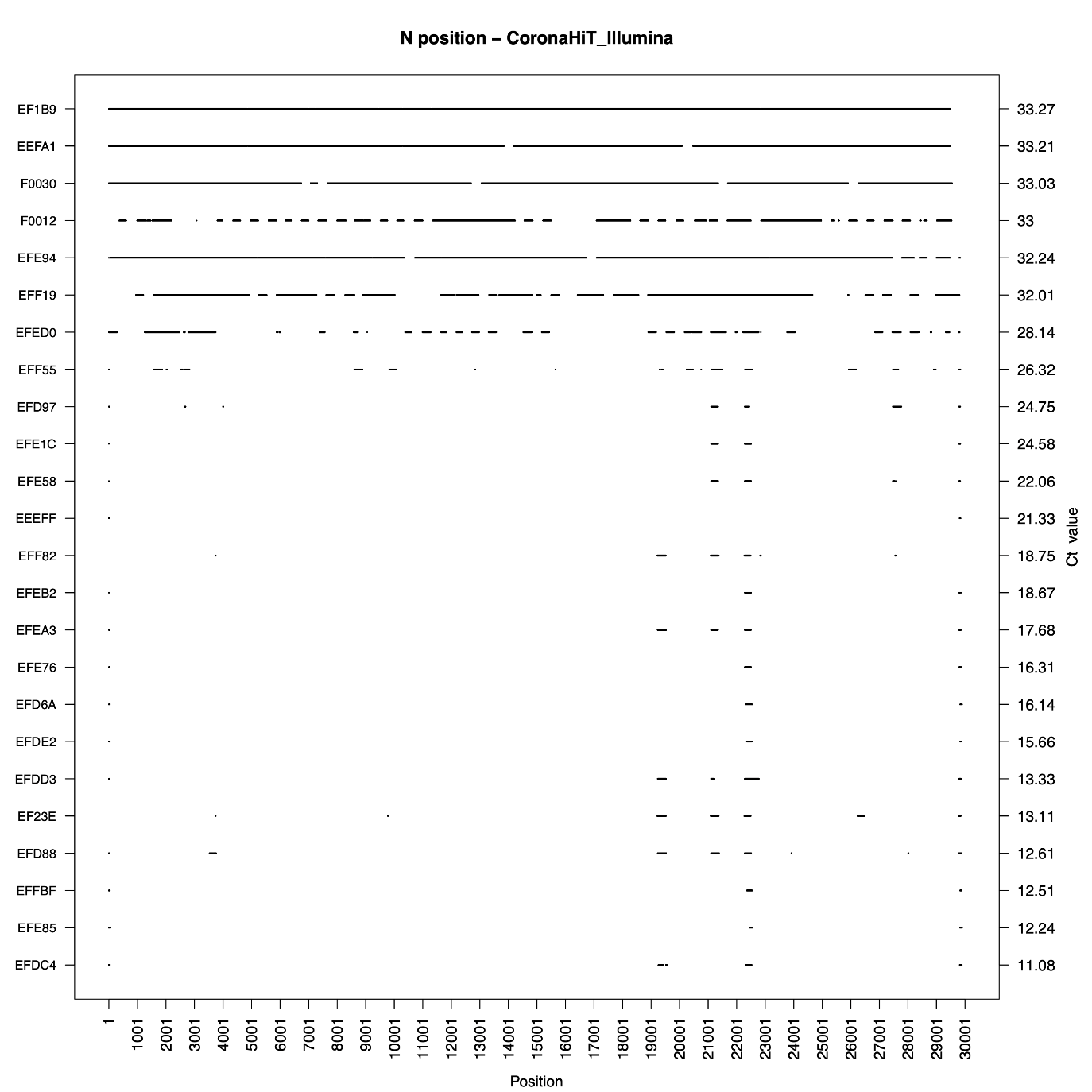

**Supplementary Figure 1**: Overview of the SARS-CoV-2 genome against the consensus sequence of each sample that had a known Ct using (a) ARTIC LoCost for routine samples, (b) CoronaHiT-ONT for routine samples, (c) CoronaHiT-Illumina for routine samples, (d) ARTIC LoCost for rapid response samples, (e) CoronaHiT-ONT for rapid response samples, (f) CoronaHiT-Illumina for rapid response samples. Black indicates a region where there is missing data, or where the coverage dropped below 20X for CoronaHit and ARTIC ONT and 10X for Illumina.
